# Across atoms to crossing continents: application of similarity measures to biological location data

**DOI:** 10.1101/2022.06.20.496870

**Authors:** Fabian Schuhmann, Leonie Ryvkin, James D. McLaren, Luca Gerhards, Ilia A. Solov’yov

## Abstract

Biological processes involve movements across all measurable scales. Similarity measures can be applied to compare and analyze these movements but differ in how differences in movement are aggregated across space and time. The present study reviews frequently-used similarity measures, such as the Hausdorff distance, Frechet distance, Dynamic Time Warping, and Longest Common Subsequence, jointly with several measures less used in biological applications (Wasserstein distance, weak Fréchet distance, and Kullback-Leibler divergence), and provides computational tools for each of them that may be used in computational biology. We illustrate the use of the selected similarity measures in diagnosing differences within two extremely contrasting sets of biological data, which, remarkably, may both be relevant for magnetic field perception by migratory birds. Specifically, we assess and discuss cryptochrome protein conformational dynamics and extreme migratory trajectories of songbirds between Alaska and Africa. We highlight how similarity measures contrast regarding computational complexity and discuss those which can be useful in noise elimination or, conversely, are sensitive to spatiotemporal scales.

## Introduction

With the advent of big data in biological systems spanning nanoscale to global networks, recent advances in heuristic, statistical, and machine learning approaches offer a variety of tools and methods to assess spatiotemporal patterns in biological applications.^1–3^ An ongoing challenge is to quantify and compare sequential spatiotemporal processes, which can involve confounding environmental and individual-based factors, as well as be affected by the frequency, accuracy, and precision of measurement.^4^ Various measures of data similarity have been applied to classify and compare individual movement trajectories in both anthropogenic and ecological applications,^5–7^ and submolecular-scale movements of proteins.^8,9^

Biological data sets inevitably contain *noise*, for example, through limits in measurement precision and accuracy, but also through actual movement not relevant to the questions of interest. For example, atoms in a protein may undergo Brownian motion at the nanometer scale, while migrating birds can forage and relocate up to hundreds of km during extended stopover periods, independently of their long-distance migratory orientation and navigation process *per se*.^10,11^ It is, therefore, natural to expect that the similarity measures of biological data will behave differently at different spatial and temporal scales.

The present study discusses known similarity measures that are potentially useful in life science applications. The similarity measures provide tools to compare two data sets (trajectories), where difference in similiarity may reflect an effect of a perturbation, deviation or variation in the studied system. For example, reduced magnitudes in similarity may reflect a conformational change of a protein or variability in flight schedules and trajectories of migratory birds. More generally, the similarity measures allow the quantification and assessment of the similarity or difference between two entities in both time and space.

The basic principle of a similarity measure is to compare two data sets that correspond to spatial or temporal variation. For example, these variations can describe the changing position of a residue in a protein or provide location information of a bird as it migrates across continents. Recent studies have highlighted the broad usability of similarity measures to distinguish among known contrasting synthetically simulated and measured trajectories.^5,6^

To demonstrate how similarity measures can be used in life sciences, we have applied the similarity measures listed in Table 1 to two distinct but related examples. The measures were applied to biological data related to magnetic field sensing in migratory birds.^12–17^ **At the microscopic level**, the cryptochrome 4 protein^18^ was suggested to be a specific receptor inside migratory birds to endow them with a magnetic compass sense. ^19,20^ Upon its biological activation, the protein is expected to change its conformation, which leads to a distinct biological function. ^13^ The change in cryptochrome dynamics upon activation calls for similarity measure analyses. ^9^ **On the macroscopic level**, the migratory orientation and movement of birds are also influenced by the Earth’s magnetic field, ^16,17^ and can also be assessed using similarity measures. ^5,6^ Differences between their individual flight trajectories can additionally be important in assessing ecological hazards, conservation concerns and anthropogenic impacts.^21,22^ For example, the migratory songbird Northern Wheatear (*Oenanthe oenanthe*) undertakes twice-yearly cross-continental journeys between Alaska and Africa.^23^ In the present paper, we apply the different similarity measures to the data sets available for both examples to showcase that such an analysis can produce either a noise-filtering or signal-inhibiting effect. For this, we have developed a python-based program package that provides straightforward usability of several similarity measures and data sets. We show that the results depend on the peculiarities of a specific example and lead to enhancing or inhibiting certain features of the analyzed data. With the performed analyses, we argue that similarity measures can serve as a powerful tool to quantify and understand spatiotemporal variability for many dynamic biological processes.

**Table 1:** Overview of selected similarity measures. The table shows the different studied similarity measures and the corresponding abbreviations. Additional information on the similarity measure is also provided and explained further in the text

| Abbreviation | Name | Attributes |  |
| --- | --- | --- | --- |
| DFD | Discrete **F**réchet **D**istance | Bottleneck | Order-dependent |
| DWFD | Discrete **W**eak **F**réchet **D**istance | Bottleneck | Order-dependent |
| DTW | **D**ynamic **T**ime **W**arping | Aggregated | Order-dependent |
| HD | **H**ausdorff **D**istance | Bottleneck | Order-independent |
| LCSS | **L**ongest **C**ommon **S**ub**S**equence | Aggregated | Order-dependent |
| DDM | **D**ifference **D**istance **M**atrix | Aggregated | Order-independent |
| WD | **W**asserstein **D**istance | Aggregated | Order-independent |
| KLD | **K**ullback-**L**eibler **D**ivergence | Aggregated | Order-independent |

## Methods

Similarity measures quantify the difference between two spatiotemporal datasets, called trajectories. A trajectory can be defined as an ordered sequence of locations. A single location within a trajectory is called an element. In the present study, all locations are considered to be in three-dimensional space. In some cases, a trajectory needs to be reduced to three probability distributions with each distribution containing information from a single spatial dimension of the three-dimensional trajectory.

Here, we introduce each studied similarity measure heuristically. Rigorous definitions, algorithmic implementations, characteristic runtimes, and detailed specific properties are given in the Supplementary Material (SM). In general, the similarity measures considered can be described as being either aggregated or bottleneck measures. Attributes for each measure are listed in Table 1. An aggregated measure is influenced by every single element of each compared trajectory. Contrasting, the bottleneck measures determine a worst-case scenario to calculate the similarity, i.e., the elements from compared trajectories which exhibit the biggest difference determine the similarity. Bottleneck measures can often be identified by their use of maxima. A second attribute that is shared by all similarity measures is their dependence on the order of the location elements within the compared trajectories. Biophysical or biological spatiotemporal trajectories will naturally be ordered by design; the distinction, however, originates from the question of whether the similarity measure disregards the given order. The order-independent similarity measures interpret the trajectories as distributions or merely as an unordered set of elements.

The discrete Fréchet distance (DFD) measures the smallest distance between two element pairs belonging to the two trajectories. The trajectories are traversed in order and all elements in both trajectories have to be considered. DFD seeks to determine the largest of the smallest distances calculated, which is then called the similarity between the two trajectories. DFD is a bottleneck, order-dependent measure.

Similarly to DFD, the discrete weak Fréchet distance (DWFD) couples pairs of elements between trajectories, but, in order to minimize the largest distances, allows successive elements to traverse backward (i.e., in reverse order). This addition allows DWFD to react to outliers or evaluate trajectories that merely zigzag around each other as more similar. DWFD is a bottleneck, order-dependent measure.

Dynamic Time Warping (DTW) is a measure that was first used in speech recognition. ^24^ Given two trajectories, DTW can be used to construct pairs of elements that maintain a consistent order within each trajectory. The collection of such pairs is called coupling, while the goal of the DTW is to minimize the sum of distances between paired elements to establish the optimal coupling. DTW is an aggregated, order-dependent measure.

The Hausdorff distance (HD) defines the similarity measure of two trajectories through the closest neighbor of elements between the two different trajectories. The maximum of all closest neighbor distances is the resulting Hausdorff distance. HD is a bottleneck, order-independent measure.

The longest common subsequence measure (LCSS) returns the number of consecutive elements that the two compared trajectories have in common. To quantify commonality, LCSS requires both a distance threshold (*ε*) and a time threshold (*δ*). Two elements in the studied trajectories are defined as common if they are found within the distance and the time threshold of one another. LCSS determines the largest number of consecutive common elements as a measure of the similarity between the two trajectories. LCSS is not a classical distance measure, as it quantifies commonness rather than the distance between two trajectories. LCSS is an aggregated, order-dependent measure.

Contrary to all the other similarity measures introduced above, the difference distance matrix (DDM) approach does not compare a pair of trajectories but rather compares structures. Given a structure encapsulating a number of locations, the pairwise distance between all these locations can be calculated and sorted in a symmetric matrix. Once the location data appears to be time-dependent, DDM calculates the time average of the pairwise distances between the locations and sorts the results into a matrix. The same distance matrix can be calculated for a second structure of equal size. At this stage, two matrices describing the distances between all elements in the two structures have been calculated. Considering the element-wise difference of the matrices quantifies the change in displacement from one set to the other. Averaging over all columns of the final matrix yields a comparable result to the other similarity measures. As DDM requires two comparable structures (sets) of trajectories, its application is more limited. The approach was, however, successfully used in earlier studies.^9,25^ DDM is an aggregated, order-independent measure.

The Wasserstein distance (WD) is a similarity measure used for comparing probability distributions. The distribution can be extracted from spatiotemporal trajectories as described above. The WD measure is the sum of distances between the two distributions. It is also known as the earth mover’s distance.^26,27^ WD is an aggregated, order-independent measure.

The Kullback-Leibler divergence (KLD) is also applied to probability distributions. It is also called relative entropy^28^ and is commonly used in machine learning.^29^ Intuitively, it quantifies how well one can distinguish between two given distributions by summing over the scaled ratio between two corresponding elements of the two distributions. KLD is an aggregated, order-independent measure.

## Implementation

To calculate and assess similarity measures, we have developed a python package SiMBols (**Si**milarity **M**easures for **B**i**ol**ogical **S**ystems) and made it available at *https://gitlab.uni-oldenburg.de/quantbiolab/simbols*. The WD, the HD, and the KLD are already available within the scipy package.^30^ DTW has been made available in the dtaidistance package.^31^ The preexisting measures were incorporated into the framework such that the same form of input data can be used for all similarity measures. We have implemented DFD, DWFD, DDM, and LCSS directly into the SiMBols package. Our implementation of DFD is optimized for memory enabling comparisons among trajectories comprised of significant amounts of elements. The implementation of DWFD employs graph algorithms supplied by the networkx package.^32^

Preprocessing routines are supplied to transfer arbitrary sequences of three-dimensional Euclidean locations into an input format understood by each measure. DTW, KLD, DWFD, DFD, and LCSS have built-in parallelization, allowing for faster computation utilizing multiple CPUs. The python package numpy^33^ was used for all described methods.

The computation time for calculating the different similarity measures can differ significantly. An account of the mathematical complexity is given in the Supporting Material (SM). In order to supply an intuition of the required computation time, all similarity measures were used to compare two sets of 497 trajectories which contained 200 elements each. Table 2 shows the benchmark times to finalize each measurement computed on a single core on Intel(R) Xeon(R) Gold 5218 CPU at 2.30 GHz.

**Table 2:** Computation time needed for similarity measure calculations. The similarity measures are sorted by the utilized computation time from the fastest to the slowest.

| Similarity measure | WD | HD | KLD | DFD | DDM | DWFD | LCSS | DTW |
| --- | --- | --- | --- | --- | --- | --- | --- | --- |
| time (sec) | 0.12 | 0.19 | 12.61 | 21.03 | 72.42 | 87.32 | 104.59 | 160.26 |

The SiMBols package was originally developed to analyse protein trajectories, before we generalized it to accept arbitrary trajectories. Therefore, SiMBols includes the possibility to read in protein simulation data and not only reduce it to the necessary geometric properties but also align and superimpose two structures of the same length^9^ which is necessary for a sensible interpretation of the similarities between protein structure simulations. The reading of protein simulation files is done using the mdtraj package,^34^ while the structural alignment employs the Kabsch Algorithm^35^ as implemented in the rmsd package.^36^ An example workflow, including the protein preprocessing tasks, is included in the SM.

## Similarity measures in protein conformational dynamics

Investigations of protein activation often focus on motion and dynamical traits. For instance, activation of a protein can induce structural changes that might initiate subsequent biophysical processes which lead to two natural questions: What are the most versatile regions within a protein structure? How is the motion influenced by external perturbations?

Similarity measures provide a tool to answer the raised questions. In earlier studies, DFD and HD have been applied to assess interprotein motions.^8^ More recently, developments and increased feasibility of all-atom molecular dynamics (MD) simulations have significantly enhanced understanding of the fundamental molecular biophysical processes.^37,38^ Evaluation of MD results often relies on the analysis of the evolution of a molecular structure and comparison of its temporal trajectory to a single reference structure through the established technique of the root mean square deviation (RMSD) analysis. Alternatively, a comparison of fluctuations within a protein structure may be performed through the root mean square fluctuation (RMSF) approach.^39^ Both of these strategies are, however, inadequate when comparing two dynamic trajectories of a protein qualitatively as both approaches only involve a comparison of structures to a reference one and not a dynamic protein trajectory.

Based on an earlier study, ^9^ similarity measures have been applied to address the questions mentioned above. Specifically, the crystal structure of pigeon cryptochrome 4 (ClCry4)^18^ was simulated dynamically in two different biological states, characterized through the redox states of the flavin-adenine-dinucleotide (FAD) cofactor and a tryptophan residue (Trp369 = TrpD).^12,13^ The redox change of these two compounds facilitates a dynamical process within the protein, leading to a structural rearrangement. This rearrangement is thought to initiate a neurological signal transfer that may be linked with the proposed role of cryptochrome in night-migratory songbirds’ magnetoreception.^40–42^ The two simulated states of ClCry4 resemble an inactive dark state (DS) and a light-activated radical pair state (RPD) specific to the redox states of the aforementioned FAD cofactor and the TrpD residue.

To test whether the activated ClCry4 would revert to its inactive DS conformation, we have performed MD simulations, where at the very beginning of the simulation the RPD conformation of ClCry4 assumed the redox states of the FAD and TrpD cofactor as exhibited in the inactive DS. In total, two replica simulations to describe the RPD→DS transition were conducted, denoted as Reverted1 and Reverted2. Additionally, two separate DS and RPD simulations were performed to establish a basis for the similarity comparison measures. A detailed description of the simulation parameters can be found in the SM.

Employing the different similarity measures, the internal conformational changes associated with the RPD→DS change in ClCry4 were probed to establish whether the rearrangements that were originally observed in the DS→RPD transition revert if the activated protein is assumed inactive.

In order to apply similarity measures, one needs to consider all temporal snapshots for each atom in the protein’s trajectory. Figure 1 illustrates how a trajectory for one atom is perceived. Every amino acid residue in a protein consists of a backbone and a sidechain. Figure 1 features an exemplary Alanine, namely the Ala230 residue from ClCry4. The backbone is the same for all residues and consists of a nitrogen atom, two carbon atoms, and an oxygen atom. The side chain is chemically attached to the carbon atom, which has a bond to the nitrogen (Fig. 1B). This carbon atom, denoted as C_α_, is thus, consistently placed at the core of each amino acid residue. Therefore, it is favorable to consider the C_α_ atom as a reference position for each residue in a protein as a well-suited focal point. Hence, the trajectory for a given residue is a sequence of a single location per time instance and should not be confused with the trajectory of the entire protein, which contains the location for all atoms in the protein.

**Figure 1:**
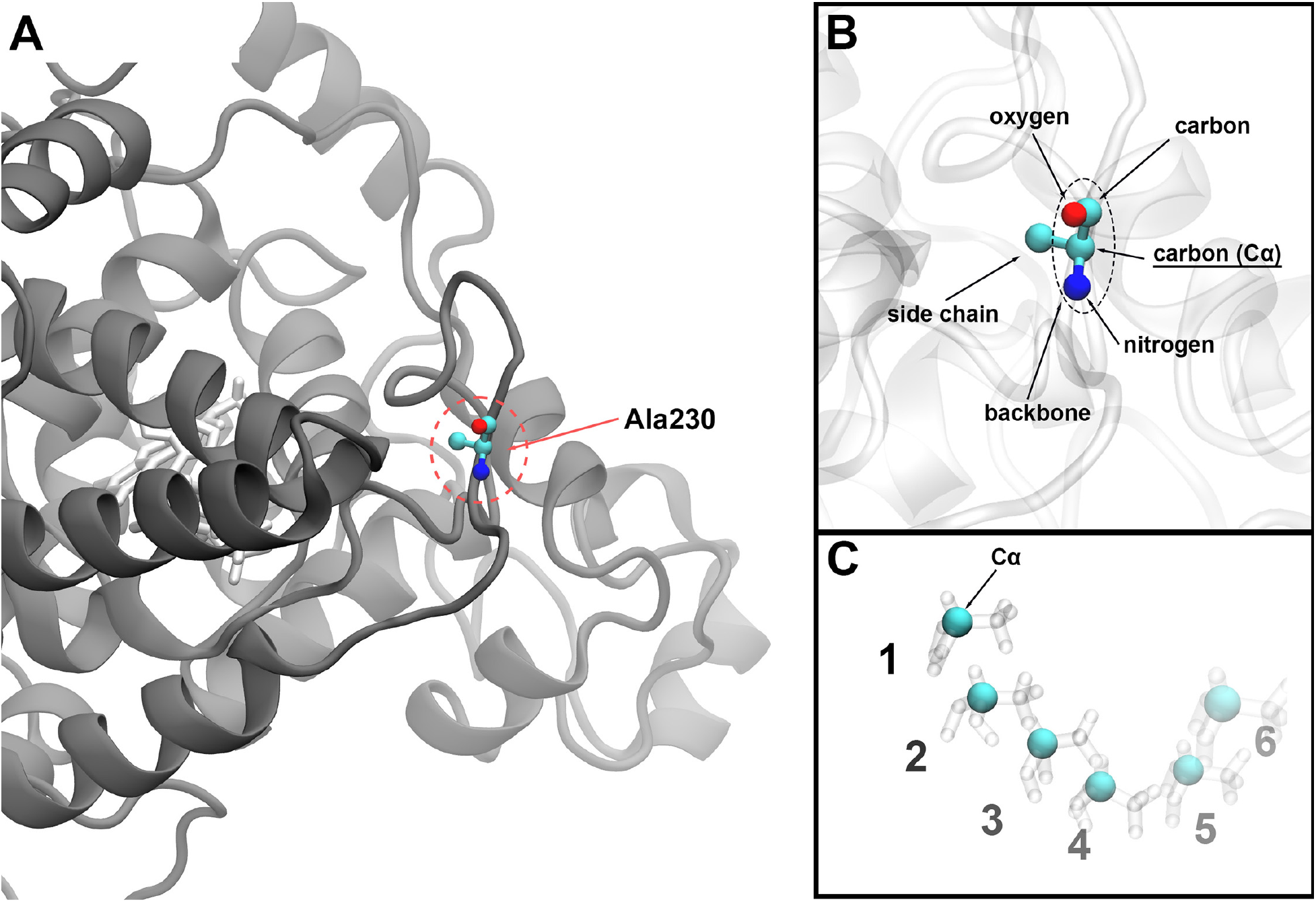
Generation of a trajectory in ClCry4. The approach for defining a simulation trajectory used as a measurable instance for similarity measures is visualized. Every residue, in this example alanine 230 of ClCry4, is individually analyzed (A). The trajectory is then considered for the respective C_α_ atom of the selected residue (B). Over time, the atom moves, yielding an ordered sequence of spatial locations. These locations then form the considered trajectory (C).

The resulting residue trajectories, extracted from the ClCry4 MD simulations, were compared using the similarity measures. Each similarity measure was computed for the respective pairs of ClCry4 states: (DS, Reverted1), (DS, Reverted2), (RPD, Reverted1), and (RPD, Reverted2).

The ClCry4 structure consists of 497 amino acid residues. Following the scheme outlined in Fig. 1, 497 individual residue trajectories were created describing the positions of each residue for all timesteps of the performed MD simulations. Every single residue trajectory has its counterpart in the to-be-compared protein simulation. Since the residues are bound to their neighbors, it is natural to expect that once a certain residue experiences a noticeable difference in the similarity measure, its neighboring residues experience some movement as well. This dependency between the individual residue trajectory pairs allows the visualization of the results for all residue trajectories simultaneously, as shown in Fig. 2. The figure shows the similarity measures computed for the comparisons of the Reverted1 simulation with both the DS (red) and the RPD (blue) state. The analysis of comparisons for the Reverted2 simulation is shown in Fig. S1 in the SM.

**Figure 2:**
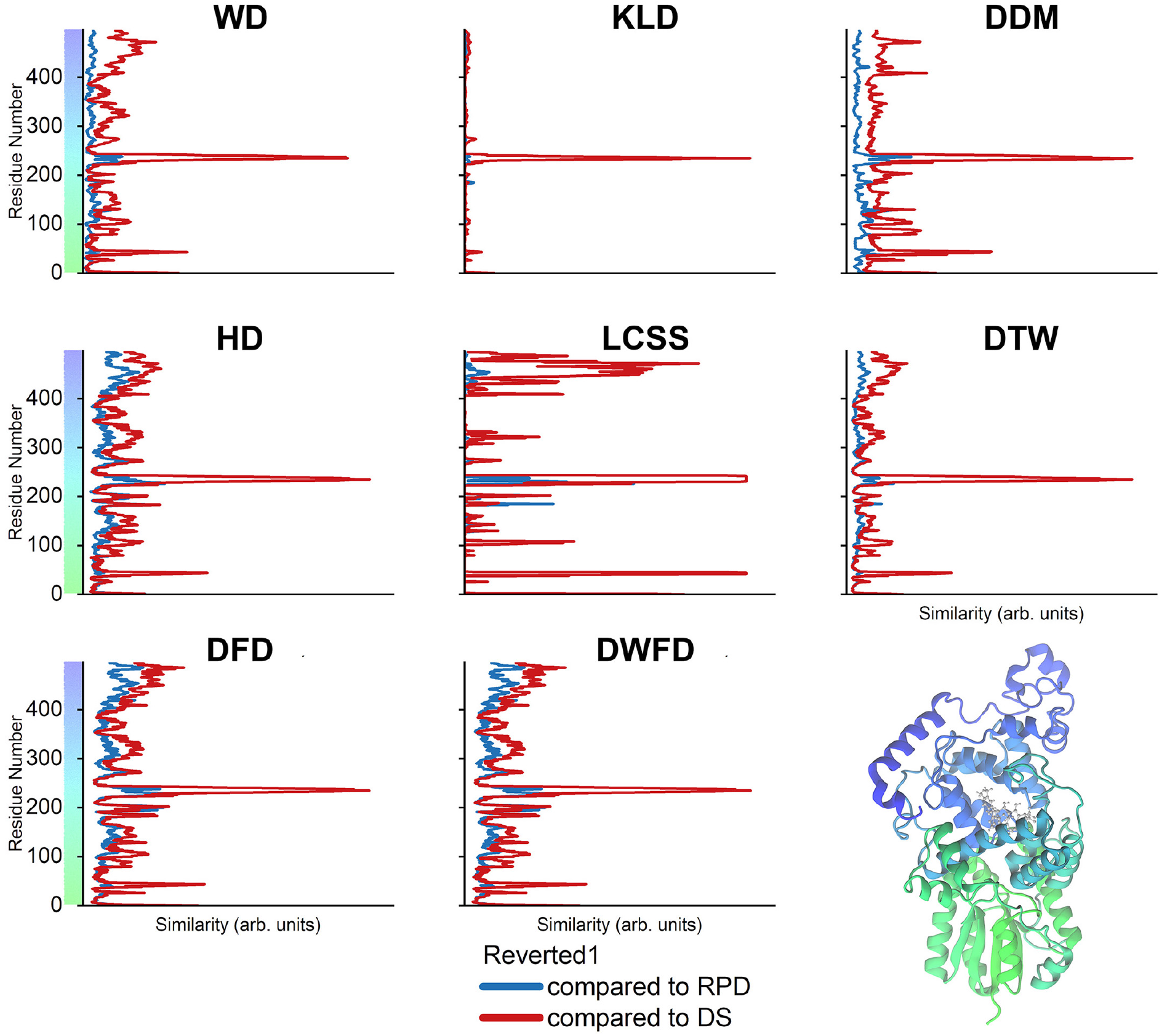
Similarity measures for ClCry4 residue trajectories. Comparison of the residue trajectories for two trajectories (RPD, Reverted1), shown in blue, and (DS, Re-vertedl), shown in red. The color scale hints towards the location of each residue in the protein structure, mapped onto its cartoon representation. Each similarity measure is computed for a trajectory of the *C_α_* atoms in each residue’s backbone. The graphs show the calculated similarity per residue. A visual comparison reveals a common peak at residues 220-240 (known as the phosphate-binding loop), which was shown to exhibit versatility in movement.^9^ The most notable differences in similarity measures can be seen in the non-prominent changes. For instance, a second peak can be observed at residues 40-50, which is hardly visible in the DFD as it becomes less significant among the noise and other fluctuations in the proteins. On the other hand, DTW distinguishes differently between noise and signal, resulting in a better distinguishable peak at residues 40-50. The results for the replica simulations comparison are shown in Fig. S1 in the SM.

All similarity measures reveal that the Reverted1 and RPD (blue) trajectories are very similar, with a notable dissimilarity between residue 220-240. These residues form the so-called phosphate-binding loop where the radical pair between FAD and TrpD is formed. On the other hand, for the comparison between Reverted1 and DS (red), more significant dissimilarities have in general been observed. Here, the dissimilarity for residue 220-240 is much greater for all employed methods. Additionally, a notable dissimilarity arises for the residues 40-50 for the majority of similarity measures. The results obtained with HD, DFD, and DWFD exhibit sensitivity to noise and vibrations within a residue (Fig. 2, red & blue). The sensitivity to noise, which results in increased fluctuations in comparison to KLD, can be explained by the fact that all three methods (HD, DFD, DWFD) are bottleneck measures. The KLD measure almost ignores small vibrations (noise) in each residue entirely but does not find the second smaller peak around residues 40-50 (Fig. 2, red), which is well shown in the plots computed using the WD or DDM measures. The result of the WD calculation exhibits a remarkably similar result to the plots obtained using the DFD and HD measures, suggesting that it might be a computationally efficient alternative to DFD and HD for resolving more pronounced motions. The LCSS plot exhibited extremes with the chosen spatial threshold *ε* = 0.5 and reveal that the phosphate-binding loop, which is near the active site cofactors FAD/TrpD, moves differently compared to other similarity measures plots. The region around residues 40-50 and the C-terminal are also significantly different compared to the other plots shown in Fig. 2. Caution needs to be exercised when working with the LCSS measure, as the spatial threshold *ε* has to be chosen manually, and the magnitude of the threshold is highly dependent on the data used. LCSS is thus highly sensitive to the choice of the parameter *ε*. Figure 3 visualizes the severe change in the similarity measurement results depending on different *ε* value. Unfortunately, there is no general guideline to determine the spatial threshold for arbitrary datasets, which is a major drawback of the LCSS similarity measure in the absence of predetermined scales of interest. During an MD simulation, a residue might not move significantly and just experience vibrations and no conformational change. In such a scenario, the start and the endpoint of the temporal trajectory are virtually similar. This peculiarity exhibited by protein trajectories results in an increased sensitivity induced by the choice of the spatial threshold *ε* because a small threshold will mark the whole trajectory as similar for an immobile residue.

**Figure 3:**
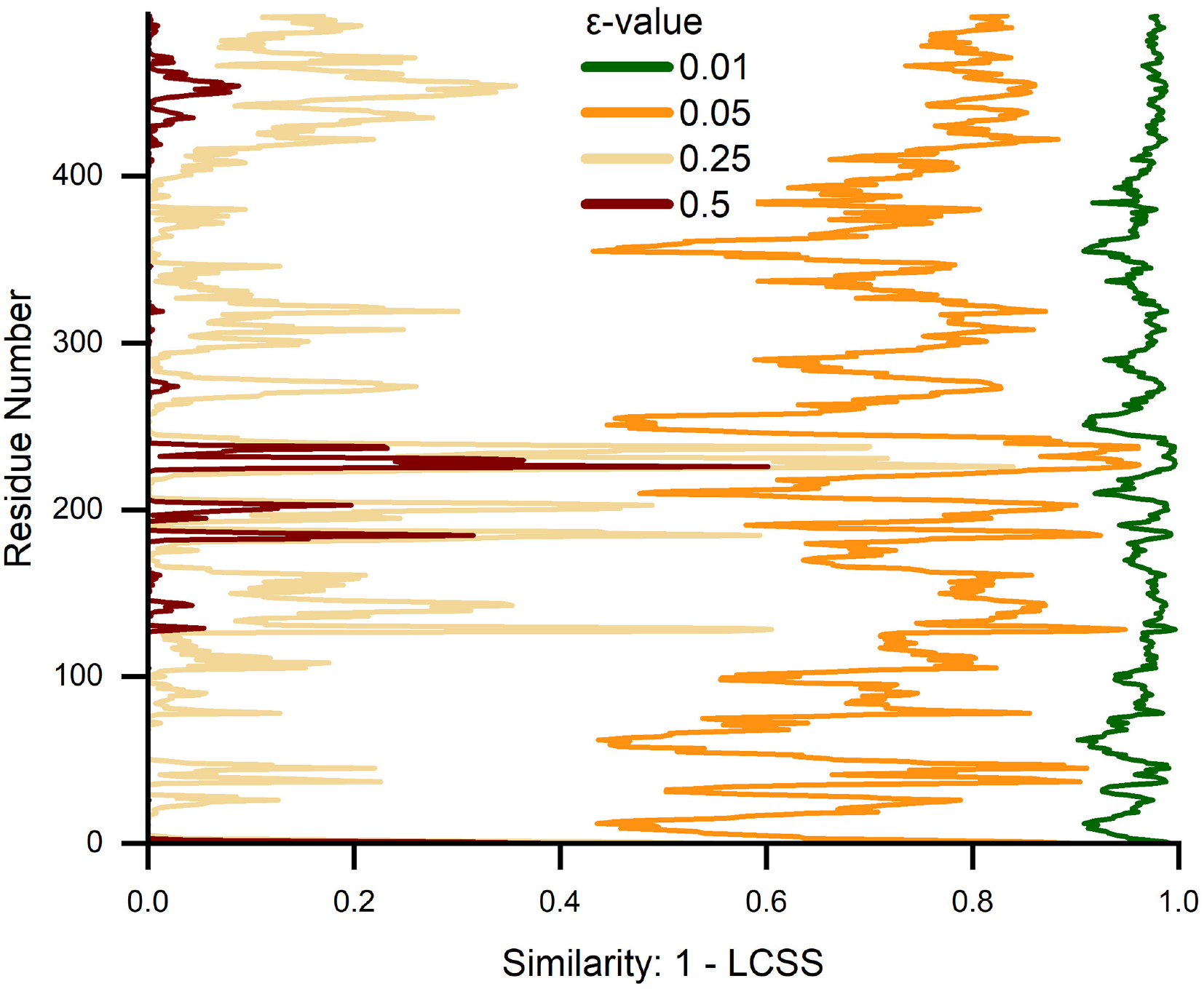
Changes in parameter *ε* lead to drastical changes in the LCSS results. The LCSS similarity for the residue trajectory comparisons for different spatial distance thresholds *ε* indicate the importance and difficulty in choosing the right *ε* threshold for LCSS, especially in protein structures, in which some trajectories might merely vibrate.

In biophysical terms, all similarity measure results show a distinct peak at the phosphate-binding loop when comparing the Reverted1 to the DS trajectory. In contrast to this, no such peak is observed for the comparison of Reverted1 with the RPD. Interpreting these differences in similarity in the phosphate-binding loop by employing results from an earlier study^9^ suggests that the conformational rearrangements observed during the activation of ClCry4 do not revert for the redox states of the FAD and TrpD cofactors and the ClCry4 structure remains locked in the established RPD conformation. A possible explanation for this behavior might be given by the solvent that was now able to reach the FAD after the phosphate-binding loop moved out of the way and is now blocking the way back, hindering the reverting motion. The effect of the solvent on the FAD might also initiate some further downstream processes or reactions, forcing the ClCry4 protein into an intermediate state.

## Similarity measures in bird flight trajectories

Macroscopic studies of migratory bird navigation behavior are confronted with a manifold of underlying factors and processes, ^43^ some relating to magnetoreception. ^16,17,44^ Some of these processes may be diagnosable using similiarty measures, e.g.: How similar are the migration routes (trajectories) of two individually traveling birds? Do birds from one specific population or age cohort have similar stopover locations and duration while migrating?

Based on location data from an earlier study,^23^ we examined the autumnal migration of eight Northern Wheatears (*Oenanthe oenanthe*, wheatear hereafter) using similarity measures. The wheatear is a night-migrant which travels twice-yearly between the Northern latitudes (Europe, Asia and North America) and Africa. Methods like GPS are not yet suitable for tracking light-bodied songbirds such as the wheatear (< 25 *g*) across their entire migratory routes. Light-level geolocation is a viable tracking method for doing so, though it only reveals twice-daily location estimates, and is less accurate than GPS. ^45^ Birds are first captured, typically near the breeding grounds, equipped with geolocators and subsequently released. Once a bird is recpatured and tag recovered, geolocator data yields two locations a day, one in the morning and one in the evening, measured at dawn and dusk. Since wheatears are night-migratory, the two locations generally do not differ a lot during the day. We, therefore, averaged the two daily locations to form one point in the flight path trajectory. However, the noise due to the geolocation method increases significantly if birds are traveling during the equinox or are close to the equator. In general, the precision of geolocation estimates is in the range of ~ 100-200 km but much less precise (~ 400 km) in the latitudinal direction when daylength becomes nearly uniform across latitudes, i.e. during periods close to the spring and autumnal equinox and near the equator. ^46^

More generally, the variability in migratory schedules among individual birds creates deviations that infer problems in the comparison among flight paths. Long-distance migratory birds typically undertake sequences of nightly flights interspersed by extended multi-day stopovers to replenish energy reserves. For the wheatears, autumn stopovers were typically clustered in Kazakhstan and last 5-20 days. ^23^ Furthermore, individual birds might not only start their journey on different days but also vary in their stopover duration.

Figure 4 illustrates location estimates during autumn migration for three of the eight individual wheatears labeled 7902, 7920, and B070. ^23^ The solid lines describe the mean migration trajectories. The colored symbols depict random daily sampled locations of each individual based on the mean and standard (normal) errors in latitude and longitude.

**Figure 4:**
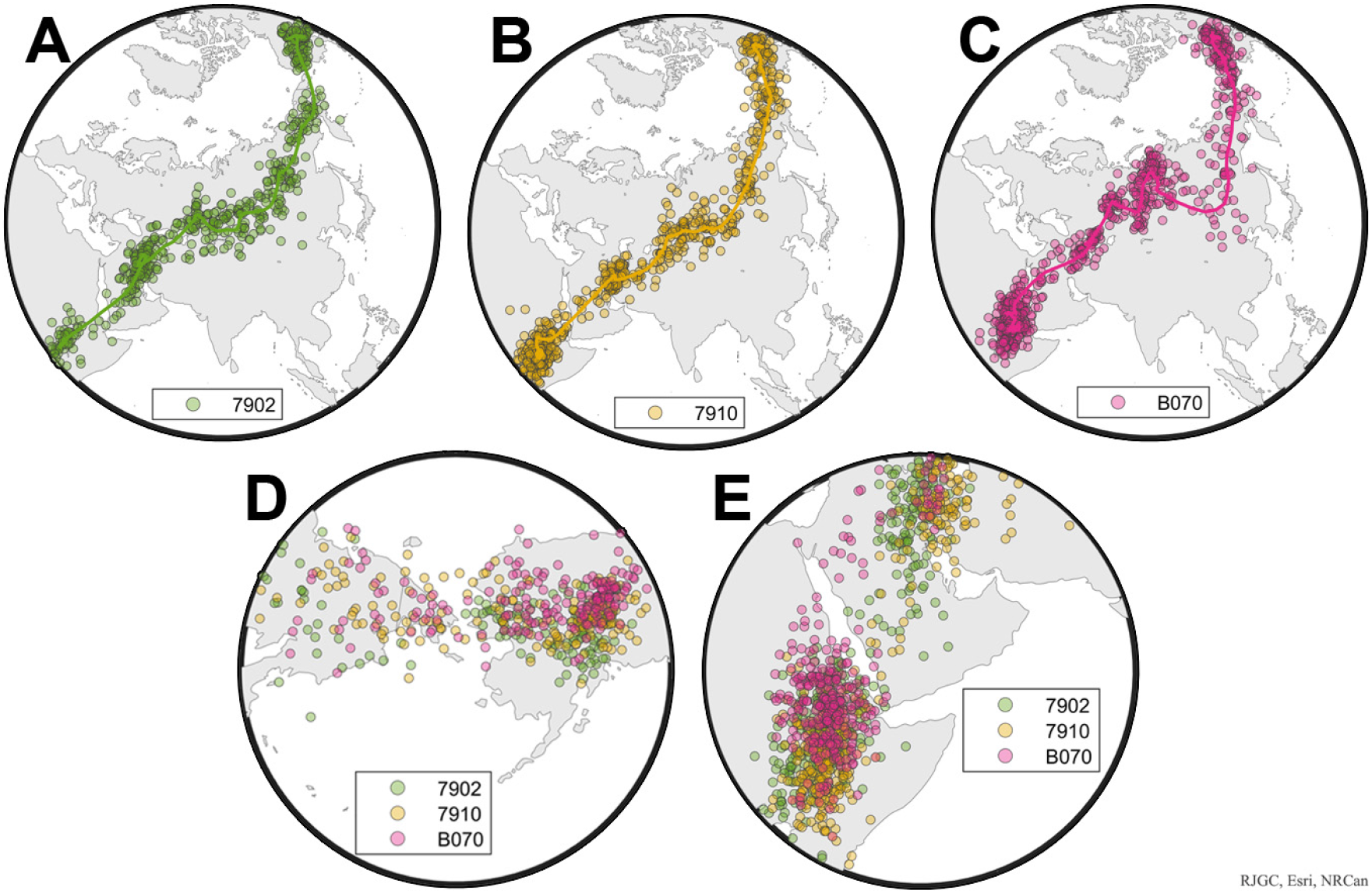
Autumn trajectories of continent-crossing songbirds. (A-C): Estimated (mean) daily locations (solid lines) with randomly sampled daily location estimates (colored symbols) based on geolocator tracking data of three Northern Wheatears (*Oenanthe oenanthe*) migrating between Alaska and East Africa, with tag labels 7902 (green line and diamonds, A), 7910 (yellow line and symbols, B) and B070 (pink line and diamonds, C); see^23^). Randomly sampled locations of all three individuals between (D) capture in Alaska and migration across East Siberia and (E) the Arabian Desert and arrival in East Africa. The map was produced with the Mapping Toolbox in MatLab 2022a. ^47^

While the data in Fig. 4A-B are visually similar, the location data of individual B070 (Fig. 4C) deviates. More precisely, B070 apparently detoured via Mongolia, while the other two individuals remained further North. Additionally, individuals 7910 and B070 exhibit closer spatiotemporal similarity in stopovers during the travel between Alaska and East Siberia (Fig. 4D) and across the Arabian Desert to East Africa (Fig. 4E) as seen by the more densely placed symbols. Trajectories of the other five recaptured wheatears are found in the SM (Fig. S2).

For the calculation of the different similarity measures, each individual bird’s trajectory was paired with all trajectories of the remaining seven birds. For clarity, we will term these comparisons as ‘‘similarity between-individuals”. Thus, 28 distinct pairs of birds’ trajectories were considered. To account for possible effects of geolocator noise on estimated similarity among trajectories of individual birds, we simulated 1,000 trajectories for each individual by sampling the daily locations including estimated deviation (for longitude and latitude) according to a normal distribution.^23^ In this way, 28,000 sampled pairs were used to assess similarity measures and a total of 8,000 pairs within an individual’s sampled trajectories. The sampling methodology and motivation are pictographicly visualized in Fig. 5. While larger samples and more advanced analysis techniques for geolocation data exist and would further reduce estimated noise, ^45^ our simple approach based on standard deviation in daily locations serves to highlight potential effects of noise on estimated similarity among measures, and is further sufficient for comparison among individual birds.

**Figure 5:**
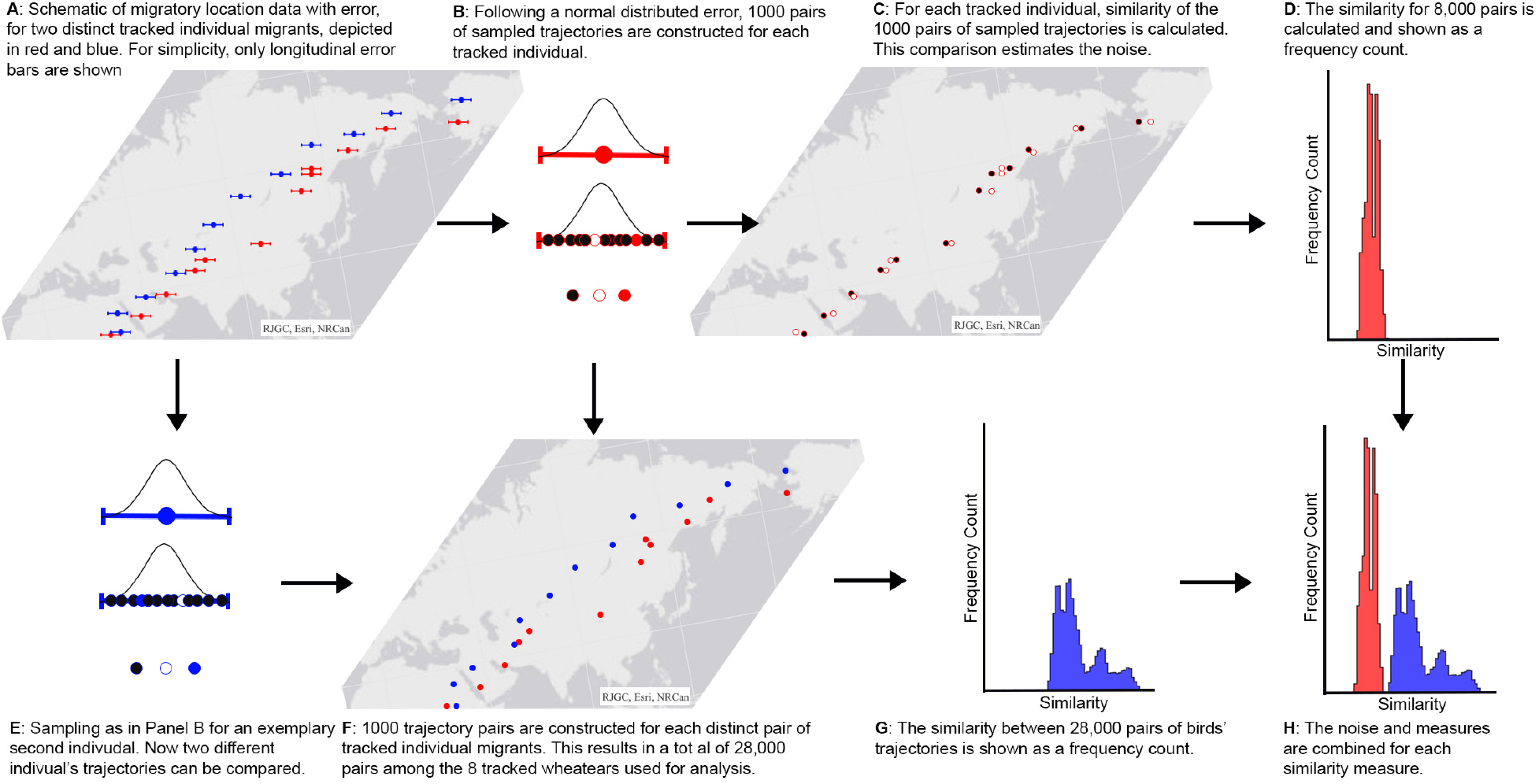
Sampling of trajectories, accounting for error. Based on location data containing error (A), trajectories are sampled for each tracked individual (B, E) according to the underlying (here, normal) distribution. Comparison between pairs of sampled trajectories from the same tracked individual (C) yields a distribution in similarity (D) to estimate noise. By comparing pairs of sampled trajectories from different tracked individuals (F), the actual similarity between two migratory trajectories can be assessed (G). A total of 28,000 comparisons were conducted. The map was produced with the Mapping Toolbox in MatLab 2022a.^47^

An intuitive comparison among measures regarding their sensitivity to deviations among trajectories can be seen in the frequency histograms of the estimated similarity among the sampled trajectory pairs (depicted in blue in Fig. 6). Note that a higher similarity value (x-axes in Fig. 6) corresponds to a greater difference between two trajectories. It is not feasible, however, to compare the absolute magnitudes of two similarity measures, as they might already differ based on their mathematical framework. To gauge how the range in sensitivity for each measure compares with the sensitivity of noise, frequency histograms of similarity between sampled trajectories from within the same individual’s sample (8,000 pairs) has been plotted in red. For comparison with the between-individual similarity (blue histograms), the within-individual values describing the noise were normalized to make the magnitudes comparable. Additionally, the LCSS measure was computed four times using different threshold values of *ε* (50 km, 100 km, 200 km, 400 km), which allows a more complete analysis while accounting for ranges within and beyond non-migratory movements during stopover periods. ^10,11^

**Figure 6:**
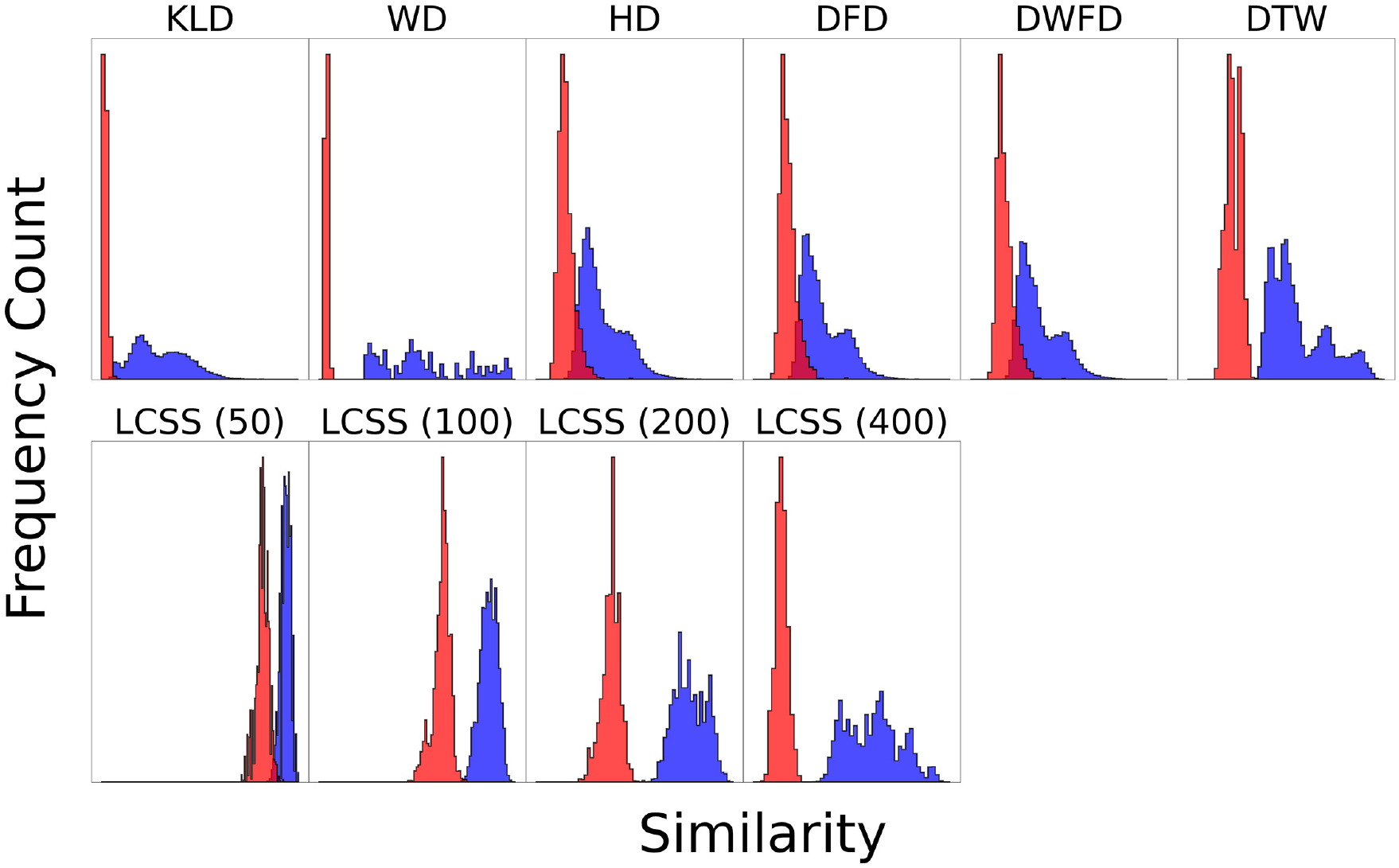
Frequency counts of similarity among migratory trajectories for each measure. Frequency count histograms for each measure are constructed (see Fig. 5) based on all 28 distinct pairs of tracked individual wheatears. Additionally, the similarity of the trajectory of one bird to itself (noise) is illustrated after being subjected to sampling (red). The red histograms, therefore, visualize the noise in the data as interpreted by the different similarity measures, while the blue histograms actually show the similarity of different trajectories. The noise histograms are normalized to make them comparable with the between-individual histograms. The greater the difference between the red and blue histograms, the better noise can be distinguished from the actual comparison. Note that HD, DFD, and DWFD have the greatest overlap between noise and similarity measure, which can be attributed to the measures being so-called bottleneck measures. DTW separates the noise and similarity measure better. KLD and WD are both insensitive to noise and therefore exhibit very narrow red histograms. LCSS reveal significant deviations for different distance threshold values (50, 100, 200, 400 km).

For KLD, a narrow noise peak can be observed while the distribution of similarity values comparing distinct individual birds is much broader. The latter is also suggestive of a bi-modal distribution. The distributions of similarity between-individuals obtained using the HD, DFD, and DWFD measures all generally resembled that of the KLD result. This is surprising due to the fact that KLD can be characterized as an aggregated method while the other three are bottleneck measures. However, KLD allows to distinguish more clearly between noise and real comparisons. In all cases, the similarity of the noise is distinguishably lower than that between different individuals. Figure 6 visualizes the difference, as the red distributions are always noticeably smaller (more left) than the blue ones.

Results obtained for the WD similarity measure show a significantly different distribution compared with the other mentioned measures. Even though WD, like KLD, is categorized as an unordered, aggregated similarity measure, and shows a similar distribution in noise, the two results reveal strong deviations. While KLD shows a bimodal-like distribution, WD shows a wide spread distribution without any clear characteristics.

For DTW, the noise can be more clearly separated from the rest of the similarity measurement as there is no overlap in distributions. Furthermore, the distribution for DTW is quadrimodal-like.

As with the protein analysis (Fig. 3), estimated similarity among the wheatear trajectories using LCSS is highly dependent on the choice of parameters. Four different spatial threshold values *ε* were employed (50 km, 100 km, 200 km, and 400 km) to demonstrate the different LCSS sensitivities. It can be observed that a higher spatial threshold leads to a higher similarity between trajectories, as can be observed in the four LCSS frequency distributions. Therefore, a meaningful choice of threshold has to be conducted carefully. For instance, a threshold of 50 km will not be sufficient to characterize whether two birds follow a similar migration route at a continental scale, ^5^ but it could be a reasonable threshold to compare foraging flights within a population at a regional scale.^5^ Calculating the LCSS for several values could conceivably facilitate differentiation between local and long-distance movements, e.g. stopover and foraging flight and directed flight.

To achieve a more detailed comparison among measures with respect to their attributes, we focus on the three trajectories depicted in Fig. 4. To facilitate comparison, we further normalized each measure to its most different trajectory pair. Again, KLD and WD show different results even though both measures can be categorized with similar attributes (aggregated and unordered). According to the KLD measure, the trajectories of the birds tagged 7910 and B070 are least similar among the three pairs, while WD indicates that the trajectories of the birds 7902 and B070 are least similar (Fig. 7). Aggregated methods are dependent with respect to every element along the trajectories and thus more sensitive to small deviations along two different trajectories. In Fig. 4 it can be observed that even though from a general perspective all three trajectories look similar, strong deviations are observed when concentrating on only few elements at the same spatial region (longitude, latitude). This deviation leads to the significant difference of the two aggregated and order-independent measures (KLD, WD). As with the distribution plots in Fig. 6, DTW, DFD, WDFD, and HD measures are in agreement and indicate that trajectory B070 differs the most when compared together with trajectories 7902 and 7910. Even though the measures differ in their type and attributes (Table 1), similar trends are observed. For this scenario, the order-dependency of DTW leads to a significant difference compared to the other two discussed aggregated measures (KLD, WD). From an intuitive perspective, the results of the DTW, DFD, WDFD and HD measures are expected when visually comparing Panels A to C in Fig. 4. Thus, to answer the the questions regarding similarity of migration routes or analyzing stopover locations of birds from the same age cohort, it might be more sufficient to use more than one similarity measure in order to interpret the used data set.

**Figure 7:**
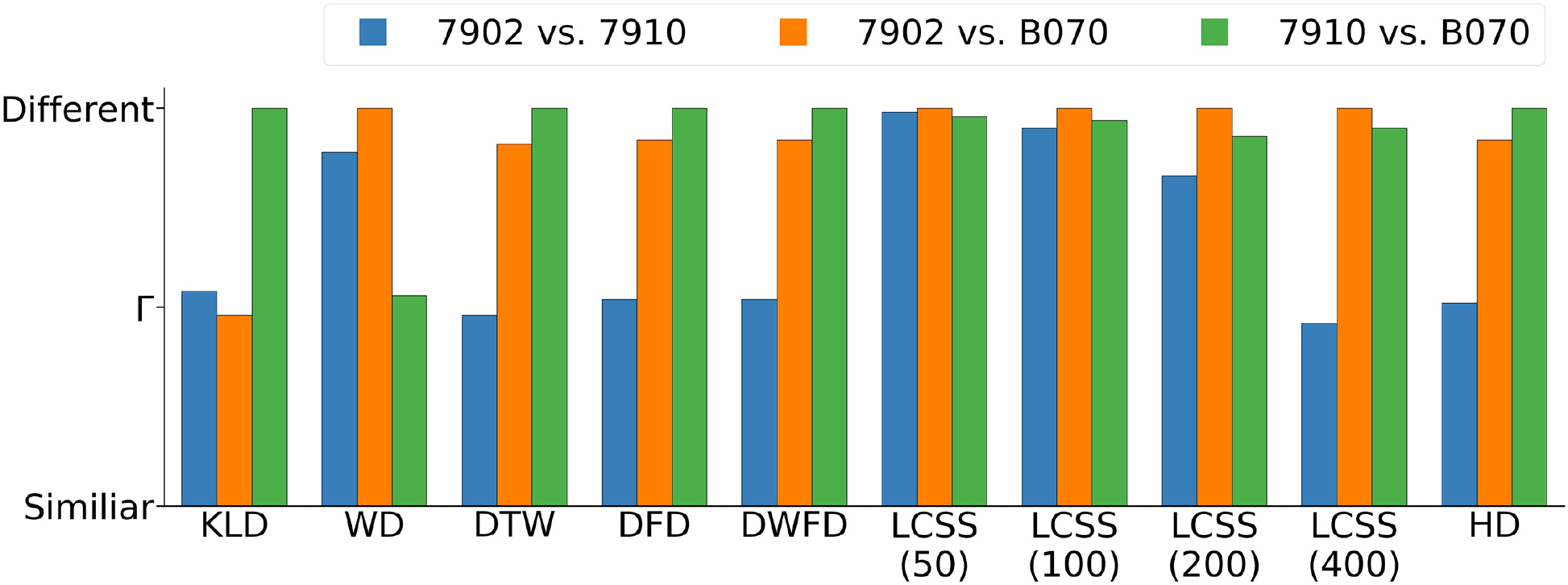
Comparison among similarity measures for three migratory trajectories. To highlight differences among similarity measures, we focused on the three tracked individual wheatear migrants depicted in Fig. 4. To facilitate this comparison, we scaled each measure to its highest dissimilarity. The symbol Γ indicates the height at which two trajectories are half as dissimilar as the most dissimilar trajectory pair. LCSS and WD both identify the trajectories of birds 7902 and B070 to be least similar, as well as WD. On the other hand, All measures except for WD indicate that individuals 7902 and 7910 followed the most similar migratory trajectory.

## Conclusion

In this work, we have presented a software package SiMBols that permits computing similarity measures for different biological systems at microscopic and macroscopic levels. SiMBols calculates the similarity measures independent of their scale, which has been demonstrated through two exemplary case studies. The package combines eight different similarity measures, which include bottleneck as well as aggregated measures. They are also distinguishable in their treatment of ordered or unordered sequences of data. Additionally, SiMBols provides a variety of preprocessing routines, which allows a versatile utilization of the package in life sciences.

We have demonstrated that SiMBols can be utilized efficiently for diverse biological problems. The extremes in the scales of the showcased case studies demonstrate that the package provides a generalized framework which can be applied to a wide variety of arbitrary spatiotemporal studies.

The similarity measures implemented in SiMBols were classified based on two major attributes, namely bottleneck or aggregated measures were discussed, while the datasets were considered being either ordered or unordered. Furthermore, it was also shown that the measures not only differ in their attributes but also in the computation time and hardware prerequisites. In general, we have demonstrated that SiMBols provides a manifold of different techniques to approach the analyses of spatiotemporal data. Even though SiMBols provides such a variety of different similarity measures, the measures for every specific study should be chosen thoughtfully. For instance, LCSS was shown to be highly dependent on the scale of the problem of interest. The magnitude of noise in the dataset was another important factor for the choice of similarity measures. It was shown that, e.g., KLD is very insensitive to noise, but might also miss smaller but yet significant differences between two trajectories.

In summary, using similarity measures provides a quantitative tool to compare and classify biological movement across scales and processes. The python package SiMBols available at *https://gitlab.uni-oldenburg.de/quantbiolab/simbols* provides a versatile computational tool to calculate different similarity measures within one unified framework.

## Acknowledgement

The authors would like to thank Jérôme Urhausen for providing valuable feedback on computation methods, coding and debugging.

Furthermore, the authors would like to thank Volkswagen Foundation (Lichtenberg Professorship to IAS), the DFG, German Research Foundation, (GRK1885 - Molecular Basis of Sensory Biology and SFB 1372 – Magnetoreception and Navigation in Vertebrates). Ministry for science and culture of Lower Saxony (Simulations meet experiments on the nanoscale: Opening up the quantum world to artificial intelligence (SMART)). Computational resources for the simulations were provided by the CARL Cluster at the Carl-von-Ossietzky Univer-sität Oldenburg, which is supported by the DFG and the ministry for science and culture of Lower Saxony. The work was also supported by the North-German Supercomputing Alliance (HLRN).

All map layers (Figures 4,5, and the figure in Appendix S2) were generated using the Matlab Toolbox^47^ using maps by Esri. The map data is the intelectual property of Esri and are used herin under liceses Copyright © Esri. All rights reserved. For more information about Esri ®, please visit www.esri.com.

## Supporting Information Available

Supplementary Material

- Similarity Measures
- Simulation methods for the protein case study
- Additional data for protein case study
- Code example for SiMBols

## Notes

https://gitlab.uni-oldenburg.de/quantbiolab/simbols

### Competing Interest Statement

The authors have declared no competing interest.

### Summary of Updates

No scientific change has been made. A mistake in the citation and acknowledging of the map material has been corrected.

