## Supplement for "Across atoms to crossing continents: application of similarity measures to biological location data"

### Similarity Measures

In this work several commonly used approaches to either compare trajectories and/or probability distributions are considered. The relevant aspects of these measures are, first and foremost, time-dependency, sensitivity to data outliers and/or noise, sensitivity to temporal or spatial changes of the data, and of course, computation time and memory consumption. Thus the methodology is subdivided into time-dependent and time-independent analyses. As time-dependent similarity measures, discrete Fréchet distance (DFD), discrete weak Fréchet distance (DWFD), dynamic time warping (DTW), and longest common subsequence (LCSS) are discussed. For time-independent approaches, the Hausdorff distance (HD), the difference distance matrix approach (DDM), as well as the Wasserstein distance (WD), and a symmetrized variant of the Kullback-Leibler divergence (KLD) are considered.

Note that the Euclidean distance serves as the underlying distance measure, i.e., the distance of two points $p, q \in \mathbb{R}^k$ is denoted as

$$d(p,q) = \sqrt{(p_1 - q_1)^2 + (p_2 - q_2)^2 + \cdots + (p_k - q_k)^2}, \tag{1}$$

where $p_i, q_i$ refer to the $i$th entries of the $k$-dimensional vectors $p$ and $q$, respectively.

The considered data sets consist of a number of timestamped point locations, so-called *fixes*, with locations given in $\mathbb{R}^2$ or $\mathbb{R}^3$. Note that it is also possible to consider polar coordinates, i.e., coordinates on a circle or a sphere specified by a radius and two or three angles. The respective transformations may change the absolute distances between two points. For example, the SiMBols package currently computes through-the-Earth distances for the bird trajectories, which differ from the great-circle distances at continental scales but do not affect relative distances arising from the comparison of point sets.

Quantifying similarity usually also involves detecting *outliers*, which are fixes that do not follow the movement pattern of their predecessors and successors. One should distinguish between *temporal outliers*, which could well follow the underlying movement pattern in terms of position but differ in time, and *spatial outliers*, where a fix is placed further away in space from other fixes which are temporally close.

The sections below discuss the different similarity measures for comparing two trajectories, that assume to have $n$ and $m$ timesteps respectively. Note that for certain measure, it might be necessary that $n = m$, as discussed below.

#### Time-dependent analyses

The following similarity measures all feature a temporal component, i.e., the timestamps are taken into consideration. For DFD and DWFD, the actual timestamps are not relevant; but the data points are given as an ordered sequence in space, whereas DTW and LCSS allow for a temporal threshold.

##### Longest Common Subsequence

The most straightforward similarity measure is the Longest Common Subsequence Measure (LCSS).

Given a temporal threshold $\delta$ and a spatial threshold $\varepsilon$, one compares two trajectories by matching fixes that have at most $\delta$ difference in time and at most $\varepsilon$ distance in space. Note that each fix of one trajectory can be matched to at most one fix of the other trajectory. The LCSS distance counts the number of consecutive fixes that do match and divides this sum by the total number of fixes. Thus, LCSS returns a value between 1 (all fixes could be matched, the curves are identical within given threshold values) and 0 (no matching fixes could be found). In contrast, larger values for other similarity measures indicate decreasing similarity between data sets. Therefore, to better compare LCSS to the other measures, it is more illustrative to analyze $1-$LCSS instead of LCSS.

Depending on the application, it can make sense to disregard $\delta$ altogether: For the prepared bird trajectories and protein dynamics analysis, the exact timing of fixes is less important than their spatial differences; thus, it is natural to set $\delta = \infty$.

As Figure 3 indicates, it can be challenging to determine a spatial threshold value $\varepsilon$ providing useful insights. Unfortunately, there is no general guideline to determine the spatial threshold for arbitrary data sets, which is a major drawback of this similarity measure in the absence of predetermined scales of interest.

Like the LCSS method itself, the choice of the spatial threshold $\varepsilon$ is highly dependent on the directedness of movement and the relevant spatial scale of biological interest in the study system. For systems characterized by undirected movement, it can become challenging to choose an appropriate $\varepsilon$, which would render the LCSS measure a less meaningful choice, as illustrated for the protein case study (see Fig. 3).

Formally, the following definitions apply:[1]

**Definition 1.** For sequences $A = \{a_1, \dots, a_n\}$ and $B = \{b_1, \dots b_m\}$ in $\mathbb{R}^d$, one defines their *longest common subsequence* as the longest sequence of strictly increasing indices $i_1, i_2, \dots i_k$ of $A$, such that $a_i = b_{i_j}$ holds for all $j = 1, \dots, k$.

LCSS is not a distance measure as such, it quantifies similarity as a proportion and consequently does not qualify as a metric, as only the symmetry condition is met. However, it is particularly useful for applications where one asks to group (parts of) data sets, as it can be easily modified to return the common subsequence of a pair of trajectories. Its sensitivity to temporal and spatial outliers depends on choices of $\delta$ and $\varepsilon$, which makes it highly adaptive to different applications, but also less intuitive to use.

From an algorithmic viewpoint, one first is required to compute a matrix with entries $d_{ij}$, where $d_{ij} = 1$ if the spatial distance of $a_i$ and $b_j$ is less than $\varepsilon$ while they also differ by at most $\delta$ steps in time. Otherwise, $d_{ij}$ is set to 0. Starting in one corner of the matrix, one should then search for a path to the opposite corner of the matrix that features the maximum number of diagonal steps. Assuming one starts at $d_{11}$, said path consists of a series of entries where each entry either lies diagonally to the lower right, directly to the right, or directly below its predecessor, such that the second entry on the path could be $d_{22}, d_{12}$ or $d_{21}$, and so on, until the element $d_{nm}$ is reached. The result is usually normalized, i.e., is divided by the smallest of the compared sequence lengths. Computation time with LCSS is dependant on the size of the matrix involved, as a constant number of computational steps per matrix entry is needed. Thus $O(nm)$ time is required to compute the LCSS of two sequences of length $n$ and $m$.

##### Fréchet/Weak-Fréchet distance

Before introducing DFD and DWFD, it is useful to first describe the classic (continuous) Fréchet distance,[2] a so-called bottleneck distance between parameterized curves in general. Fréchet distance has most often been applied to compare, characterize and cluster real-world trajectory data.[3,4]

The following analogy is often used to describe the Fréchet distance to gain intuition. Consider a person walking their dog. Each of them follows a predefined path, and they need to start and finish their walk at the same time. The Fréchet distance equals the maximum distance between the human and the dog during their walk, provided they choose their respective speeds to minimize the maximum distance, i.e., it equals the length of the shortest possible dog leash.

More formally, one considers two sets of time stamped points, e.g., geolocation data, $R = \{(p_1^r, t_1^r), \ldots, (p_n^r, t_n^r)\}$ and $S = \{(p_1^s, t_1^s), \ldots, (p_m^s, t_m^s)\}$, where $p_i$ represents a position in one, two or three dimensions, and $t_i$ are the time stamps. The points are ordered in an ascending order of timestamps. As timestamps are no longer important as soon as the data sequences are ordered, the corresponding labels can be omitted. To obtain parameterized curves, interpolation between consecutive points (fixes), should be accomplished by introducing straight line segments $\overline{p_i, p_{i+1}}$ to obtain a connected curve for both $R$ and $S$.

In the course of the Fréchet distance calculation, both curves have to be traversed in order, i.e., starting from the first fixes of the two sequences, one travels along the interpolated edges to the respective subsequent point, and so on, until both traversals reach their last fixes. The Fréchet distance equals the largest distance between a pair of fixes traversed in the process, provided the fixes are chosen to try to minimize their maximum separation.

Finally, the weak Fréchet distance, as opposed to the Fréchet distance, allows a change in direction while traversing. Thus, at times a fix might be considered which would have been impossible in the Fréchet distance calculation.

##### Discrete Fréchet Distance

Biological data almost inevitably consists of discrete sets of locations marked with corresponding time stamps. The following variant of the Fréchet distance[5] is therefore particularly useful.

**Definition 2.** The *discrete Fréchet distance*, also known as the coupling distance, of two data sets $A = \{a_1, a_2, \dots a_n\}$ and $B = \{b_1, \dots, b_m\}$, $A, B \subset \mathbb{R}^d$ is defined as

$$\delta_{DFD}(A, B) = \min_{L} \max_{i} d(a_{u_i} - b_{v_i}), \tag{2}$$

where $L$ is a *coupling*, which is defined as a sequence of point pairs $(a_{u_1}, b_{v_1}), \dots (a_{u_k}, b_{v_k})$. For these pairs, it holds that $u_1 = v_1 = 1, u_k = n, v_k = m$, while for the all intermediate $u_i$ (and analogously $v_i$) $u_{i+1} = u_i$ (or $u_{i+1} = u_i + 1$). A pair of points $a_\ell$ and $b_j$ is referred to as *matched*, if there exists $u_i, v_i$ such that $a_\ell = a_{u_i}, b_j = b_{v_i}$ and $(a_{u_i}, b_{v_i}) \in L$.

To gain intuition on the discrete Fréchet distance, imagine pebbles on a pond. Instead of continious paths, one is now tasked with crossing the pond by hopping from one pebble to the next one. The timestamps give a traversal order of the pebbles. Instead of a person and a dog, picture two frogs, each hopping on a set of pebbles. Both frogs need to travel in one direction (i.e., cross the pond) but can progress either simultaneously or one at a time. The sequence in which they progress is encoded by the coupling $L$. The question is to determine the coupling that minimizes the maximum distance between the two frogs.

The DFD is a metric, which means that it is a well defined distance measure. Let $m$ and $n$ be the number of timesteps in the two considered trajectories. Algorithmically speaking, one is required to compute (a part of) the distance matrix $D = \{d_{ij}\}$, where $d_{ij} = d^2(a_i - b_j)$ with $a_i$ and $b_j$ being the $i$th element of the first trajectory and the $j$th element of the second trajectory. Selecting $d_{11}$ in the upper left corner of the matrix, one should then determine the best path through $D$ to connect $d_{11}$ to $d_{nm}$ in the bottom right corner. Starting at $d_{11}$, this path is formed by selecting either $d_{12}, d_{21}$ or $d_{22}$, i.e., permitting navigation to the right, downwards or diagonally to the bottom right, until $d_{nm}$ is reached. Thus, the path encodes the coupling $L$. Theobjective is to find the path whose maximum entry value is as small as possible. The asymptotic runtime of the algorithm is determined by the cost of computing the distance matrix. Thus $O(nm)$ time is needed to compute the DFD of two trajectories featuring $n$ and $m$ timesteps, respectively.

**Discrete Weak Fréchet Distance**

Analogously to the Fréchet distance, DWFD allows backtracking. In the example above, the frogs are now not only allowed to hop to the next pebble but could also visit the previous pebble again. The difference in the formal definition is subtle, though:

**Definition 3.** The *discrete weak Fréchet distance* of two data sets $A = \{a_1, \dots a_n\}$ and $B = \{b_1, \dots, b_m\}$, $A, B \subset \mathbb{R}^d$ is defined as

$$\delta_{DWFD}(A, B) = \min_{L} \max_{i} d(a_{u_i} - b_{v_i}), \tag{3}$$

where $L$ is now a *semi-coupling*, which is a sequence of point pairs $(a_{u_1}, b_{v_1}), \dots (a_{u_k}, b_{v_k})$. For the pairs it now holds that $u_1 = v_1 = 1, u_k = n, v_k = m$, while for all intermediate $u_i$ (and analogously $v_i$), $u_{i+1} = u_i$ or $u_{i+1} = u_i \pm 1$.

This seemingly minor difference has major implications for the similarity measure. First, DWFD is not a metric, as two sequences $A, B$ may be distinct but still have $\delta_{DWFD}(A, B) = 0$. On the other hand, outliers with respect to the time component hardly influence the similarity results since backtracking is allowed.

DWFD is computed by means of finding an optimal path in the distance matrix $D$, where it is possible to traverse upwards and to the left, i.e., considering an entry $d_{ij}$ on the path, the subsequent entries may be $d_{i\pm1,j}, d_{i,j\pm1}$ or $d_{i+1,j+1}$. Even though the asymptotic runtime of $O(nm)$ is not affected by the change in algorithm, computation time increases significantly compared to DFD.

##### Dynamic Time Warping

DTW is a technique originally designed for speech recognition,[6] as it aligns similar patterns without disregarding the order in which they occur.

The technique is closely related to DFD; for two sequences $A, B$ of timestamped fixes, i.e., locations in $\mathbb{R}^2$ or $\mathbb{R}^3$, one seeks to find a coupling of points. However, unlike DFD, DTW is an aggregated rather than bottleneck measure. For DTW, a coupling that minimizes the sum of squared distances of all distance-minimizing pairs of locations is selected. More formally:[7]

**Definition 4.** For sequences $A = \{a_1, \ldots, a_n\}$ and $B = \{b_1, \ldots b_m\}$ in $\mathbb{R}^d$, the *dynamic time warping distance* is defined as

$$\delta_{DTW}(A, B) = \begin{cases} 0 & \text{if } A = B = \emptyset \\ \infty & \text{if } A = \emptyset \text{ or } B = \emptyset,\ A \neq B \\ d^2(a_1, b_1)) + \min\{\delta_{DTW}(A_{[2,n]}, B_{[2,m]}), & \\ \delta_{DTW}(A, B_{[2,m]}]), \delta_{DTW}(A_{[2,n]}, B)\} & \text{otherwise.} \end{cases} \tag{4}$$

Unlike DFD, DTW is not a metric as it does not obey the triangle inequality. Since the sum of distances between matched points is compared, rather than the maximum distance of all matched pairs of points, single outliers become less significant for DTW, as compared to the DFD calculation. Compared to DWFD, DTW picks up on all types of outliers, not just the spatial ones.

DTW can be computed by means of the distance matrix $D$. In that case, one should determine the path corresponding to a coupling $L^* = \min_L \sum_{(a_i,b_i)\in L} d^2(a_i, b_i)$. Note that one can additionally introduce a parameter (called $\delta$) that would only allow an "offset" of $\delta$ positions in the coupling to be considered, i.e., only allowing $u_i, v_i$ with $d(u_i, v_i) \leq \delta$. We set $\delta = \infty$ for the illustrated case studies in the main manuscript and set it as a default value in the SiMBols package.

#### Time-independent analyses

Some data sets either do not feature a temporal component at all, or their timing is not considered necessary for the specific application. The following sections therefore focus on spatial similarity only.

Specifically, the Hausdorff distance (HD), a well-known distance and similarity measure often used in shape matching,[8] the Difference Distance Matrix approach (DDM), which is used in protein analysis,[9,10] as well as Wasserstein distance (WD) and Kullback-Leibler divergence (KLD) are discussed. The latter two are similarity measures for probability distributions and are particularly useful when analyzing large data sets.

##### Hausdorff distance

The Hausdorff distance is a classic distance and similarity measure used to compare and analyze geometric shapes.[11] It is related to the Fréchet distance but does not take traversal orders into account. The Hausdorff distance can be illustrated as follows: Consider two groups of people standing at fixed positions, the first group wears red clothes and the second one blue clothes. All red-clothed people identify the distance to the closest person wearing blue and vice versa. The maximum of all distances to the closest person wearing the respective other color then equals the Hausdorff distance between the two groups.

**Definition 5.** For two point sets $P = \{p_1, \ldots, p_n\}$ and $Q = \{q_1, \ldots q_m\}$ in $\mathbb{R}^d$, the *Hausdorff* *distance* is defined as

$$\delta_H(P, Q) = \max\{\max_{p_i \in P} \min_{q_j \in Q} d(p_i, q_j), \max_{q_j \in Q} \min_{p_i \in P} d(q_j, p_i)\}. \tag{5}$$

The Hausdorff distance is a metric, and it is insensitive to temporal outliers as it does not feature a temporal component. Spatial outliers, however, are as easily picked up as with DFD; both are bottleneck distance measures.

Computation-wise, finding the closest point in set $P$ for each point in $Q$ and vice versa takes $O(nm)$ time. Determining the largest distance among the computed ones can be done without further increasing the asymptotic runtime.

##### Difference Distance Matrix approach

The following approach returns a vector of distances, where each entry encodes the distance, or rather the similarity, of one point in a point set to all points of a second point set. Consequently, the difference distance matrix approach is commonly used in protein analysis, where a residue in a protein is compared to all residues of a reference structure or the second protein.

For the difference distance matrix approach (DDM), one first is required to compute a matrix holding the normalized pairwise distances between two points of the same set. Having computed the distance matrices for the two datasets, one computes the differences between their entries and build the mean value of all entries in each column. The result is a vector holding said column sums.

**Definition 6.** Consider two sets of locations for points $P = \{\{p_1^1, \ldots, p_1^n\}, \ldots, \{p_k^1, \ldots, p_k^n\}\}$ and $Q = \{\{q_1^1, \ldots q_1^m\}, \ldots, \{q_k^1, \ldots, q_k^m\}\}$, where $p_i^\ell$ (resp. $q_j^\ell$) denote the position of point $p_i$ (resp. $q_j$) at time $\ell$. The difference distance matrix $D'$ with entries $d'_{ij}$ for one point set is then defined as

$$d'_{ij}(P) = \frac{1}{n}\sum_{\ell=1}^{n} d(p_i^\ell, p_j^\ell)$$
$$d'_{ij}(Q) = \frac{1}{m}\sum_{\ell=1}^{m} d(q_i^\ell, q_j^\ell).$$

The corresponding distances between two point sets are then given by

$$d_{DDM}(P, Q) = \frac{1}{n}\sum_{j=1}^{n} |d'_{ij}(P) - d'_{ij}(Q)|. \tag{6}$$

Note again that $d_{DDM}$ comes as an $n$-dimensional vector, where the $i$th entry encodes the distance of point $p_i \in P$ to the point set $Q$ (or vice versa for interchanged roles of $i$ and $j$, or $P$ and $Q$).

This similarity measure fulfills all requirements of a metric. The DDM method is used in protein analysis because it is remarkably robust to noise due to several normalization steps during its computation.[9,10] Single outliers are still detectable since one obtains the difference between a single point of one set to all points of the other set. Thus, it is possible to compare the distance values for all individual points of a set.

As with most similarity measures before, computation time is dominated by computing the distance matrices. There are $n + m + 1$ matrices of size $k^2$ to compute, one per time step, so the overall asymptotic runtime sums up to $O((n + m)k^2)$.

##### Wasserstein distance

The Wasserstein distance (WD) is a distance measure between normalized distributions. Thus the point sets should be treated as distributions in $U, V \subset \mathbb{R}^d$. Again, the order of points is disregarded, therefore movement trajectories are less suited for this approach. For proteins and other biochemical structures, however, the Wasserstein distance is a useful mean for analyzing similarity.[12] It is often referred to as the *Earth mover's distance* due to the following common analogy: Assume two piles of earth describe the given distributions. The Wasserstein distance equals the amount of work a person has to apply to shift the first pile to match the shape and position of the second pile.

**Definition 7.** The *Wasserstein distance* of two normalized distributions $U = \{u_1, \ldots, u_n\}$ and $V = \{v_1, \ldots, v_n\}$ in $\mathbb{R}$ is defined as

$$\delta_{WD}(U, V) = \sum_{i=0}^{n} |WD_i|, \tag{7}$$

where $WD_0 = 0$ and $WD_{i+1} = u_i + WD_i - v_i$, $i = 0, \ldots, n-1$ keep track of how much "earth" needs to be moved between consecutive points in the distribution.

The Wasserstein distance is a metric for normalized distributions. Two trajectories can only be analyzed using WD if they have the same number of timesteps, $n = m$. Consider two sets of points in $R^3$, namely the $x$, $y$ and $z$ coordinates for all elements in the trajectories. These sets are analyzed separately: First, the $x$-coordinates of the two point sets are considered as distributions and their corresponding WD is computed. This procedure is repeated for the remaining $y$- (and $z$-) coordinates and all separate WD-values are added to obtain the total distance between the point sets. Temporal outliers are only relevant if the timestamps are considered as additional coordinates. Single spatial outliers are hard to detect since the pointwise distances are summed up, while noise may increase the distance value.

The WD similarity measure is particularly fast to compute, as a constant number of operations per point is performed. In total, one needs $O(n)$ time to compute the Wasserstein distance of two sets of $n$ points each. Note that the general definition of WD for points in $\mathbb{R}^d$ for $d > 1$ is slightly more complicated, and its computation takes $O(n^3)$ time by applying the so-called Hungarian algorithm.[13]

##### Kullback–Leibler divergence

The Kullback-Leibler divergence, also called relative entropy, is an asymmetric distance between a probability distribution and its reference distribution[14]. Informally, it quantifies the information loss that arises from using $V$ as an approximation for the actual distribution $U$.

**Definition 8.** For two distributions $U, V$, as defined for the Wasserstein distance, the *Kullback-Leibler divergence* is defined as

$$d_{KLD}(U, V) = \sum_{i=1}^{n} u_i \ln\left(\frac{u_i}{v_i}\right). \tag{8}$$

As a symmetric measure is desirable to compare two sets of equally obtained data, we employ a smoothed, symmetric version:

$$\delta_{KLD}(U, V) = \frac{d_{KLD}(U, V) + d_{KLD}(V, U)}{2}. \tag{9}$$

Intuitively, KLD quantifies how well one can distinguish between the two given distributions. The Kullback-Leibler divergence itself is not a distance metric due to its asymmetric nature. The definition of KLD in Eq. (9) resolves the asymmetry. However, the triangle inequality does not hold, so KLD still does not qualify as a distance metric. Note that all values in the considered data sets must be strictly positive, which is not necessarily the case for protein data, for example. Therefore, before applying KLD, all points in the compared data sets are translated such that the minimum coordinate value for any point equals one. This measure is highly aggregated, which makes it tolerant to noise.

Computing the KLD requires computing the natural logarithm for all entries of the two data sets. As both data sets need to have the same size, one requires to compute $O(n)$ logarithms, which can then be weighted and added up in constant time. Thus $O(n)$ time to compute $\delta_{KLD}(U, V)$ is needed, assuming that the distributions $U$ and $V$ feature $n$ strictly positive entries, each.

### Simulation methods for the protein case study

All simulations were set up using the online platform VIKING.[15] Parameters were kept identical to the earlier study,[9] in which the CHARMM36 force field with CMAP corrections was used to describe the interatomic interactions.[16–23] The parameters for FAD and $FAD^{\bullet-}$ were adopted from a study by Xu *et al.*[24] The simulations were conducted using NAMD[25,26] in an NPT (constant number of particles, pressure, and temperature) statistical ensemble at a temperature of 310 K and an atmospheric pressure of 1 bar.

All simulations were conducted for 200 ns. Establishing a base for comparison the original inactive DS and the RPD state were prolongated without any change to their respective redox charges. Two replica simulations were initiated starting from the original RPD conformation, in which the charges were reverted to the inactive DS form (RPD$\rightarrow$DS).

The molecular dynamics simulation trajectories delivered the positions of all atoms in the ClCry4 protein for each snapshot in time. In order to remove global rotation and translation motions, the protein structures were aligned in an iterative process to minimize the RMSD: The whole structure of the protein was aligned to its first frame using Kabsch-Algorithm.[27] Subsequently, the structures are aligned again to the average structure after the first alignment. After each step, the RMSD was measured. The structure was considered aligned once the RMSD value reached a threshold.

### Additional data for protein case study

A replica simulation shows the very same behavior as the simulation discussed in the main text. In all the measures, the observed peaks are still in favor of the RPD state showing a locking conformation and no reverting to the original DS conformation, see Fig. S1.

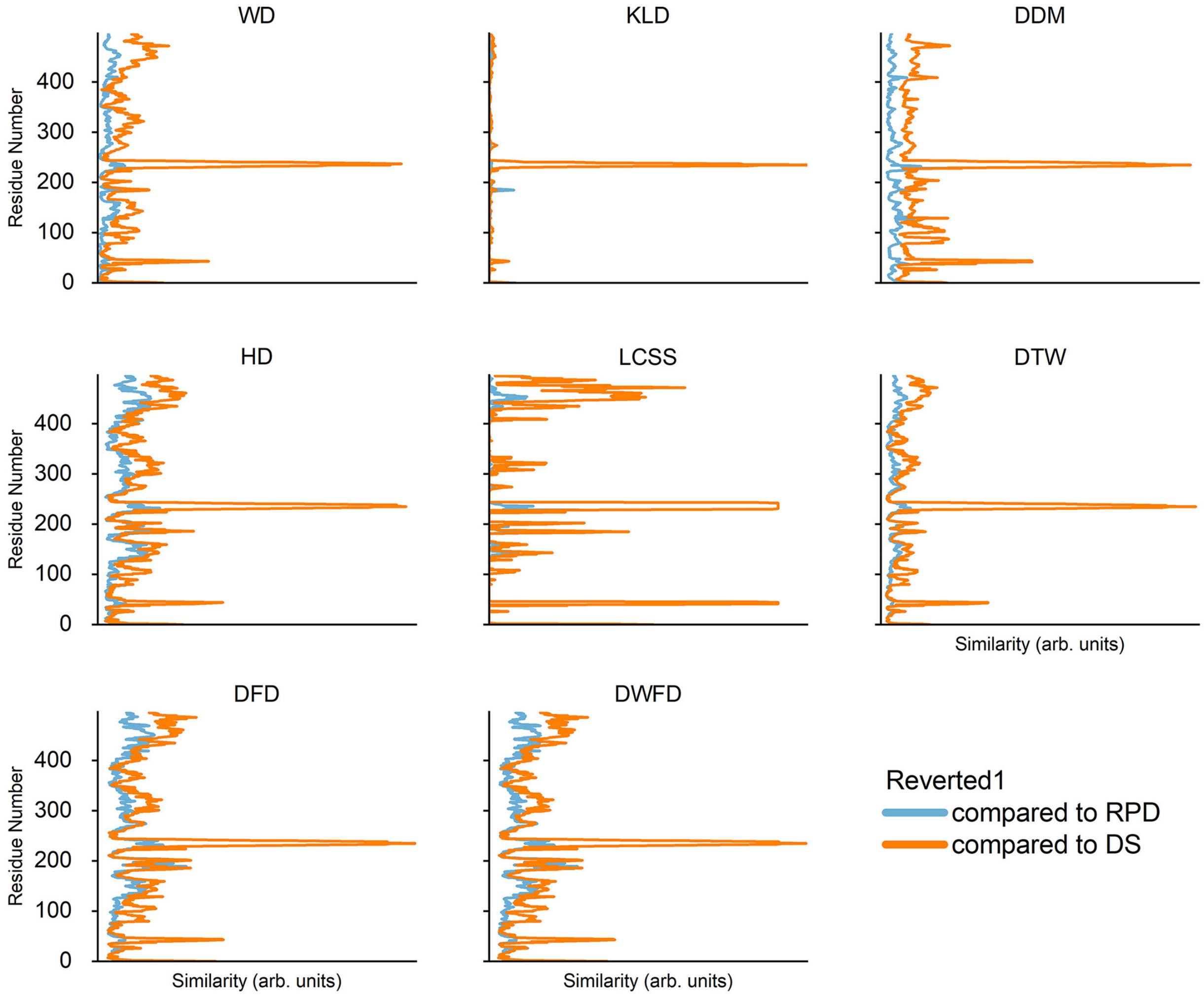


Figure S1: **Similarity Measures for Pigeon Cryptochrome 4 - Replica Simulation.** As already visualized in Fig. 2, the replica simulation shows the peak for each discussed similarity measure at residues 220 to 245. The ups and downs for each measure behave analogously as for the simulation trajectories discussed in the main paper.

### Additional bird trajectories

Figure S2 visualizes the five bird trajectories and their respective errors omitted in Fig. 4 in the main paper.

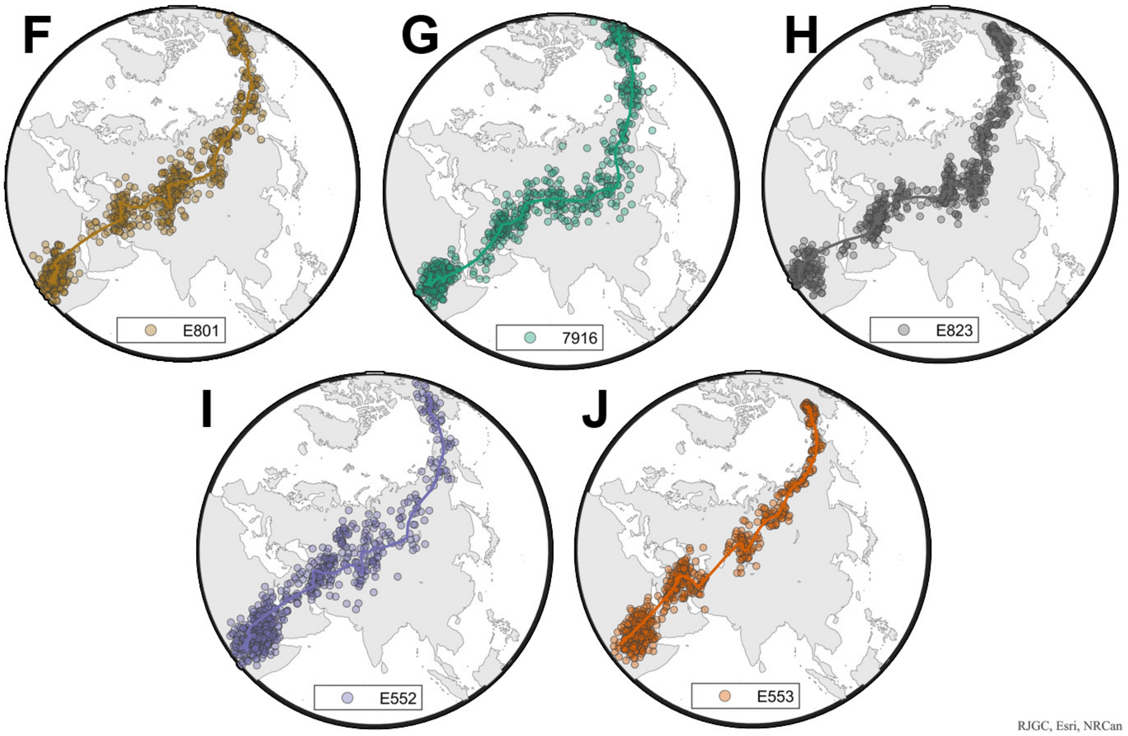


Figure S2: **The remaining 5 bird flight trajectories supplementing Fig. 4 in the main paper.** Panels A to E show the remaining 5 out of 8 bird trajectories. The map was produced with the Mapping Toolbox in MatLab 2022a.[28]

#### Code Example for SiMBols

In order to supply some guideline on how the python package SiMBols and the preprocesses for protein data can be utilized, a commented ready-to-run example workflow is provided. Needed input data, as well as this example, can be downloaded and tested at *https://gitlab.uni-oldenburg.de/quantbiolab/simbols*.

```
import SiMBols

"""We show a short example through all the steps for two protein
   trajectories. For this case, two example trajectories of Pigeon
   Cryptochrome 4 in a so-called Darkstate and a Radical Pair D state were
    truncated to just 200 snapshots. This allows for a fast exemplary
   computation. The necessary namd simulation files (.psf and .dcd) files
   are also provided to allow a trial run of the methods provided in the
```

```python
    package. We also measure the time, the example output shows the time it took on Intel(R) Core(TM) i5-8250U CPU @ 1.60GHz. It will be running on one CPU only, unless specified"""

print("Begin Example Output:")
import time

start = time.perf_counter()  # Set a start time to measure how long it takes

"""Preparation: Preparation reads a .psf and a .dcd file to transfer the location data of all the atoms to a numpy array. It takes a file location as string for the .psf, and a list of file locations as string for the dcd. This allows to load multiple .dcds. Note however, also a single .dcd needs to be supplied in a list with one element."""

"""We give the necessary paths:"""
traj1_psf = 'trajectory1.psf'  # This is the dark state trajectory psf.
traj1_dcd = ['trajectory1.dcd']  # This is the dark state trajectory dcd.
traj2_psf = 'trajectory2.psf'  # This is the radical pair D state trajectory psf.
traj2_dcd = ['trajectory2.dcd']  # This is the radical pair D state trajectory dcd.

"""Preparation.getList() returns a numpy array of shape (timesteps, atoms, 3) containing the location of each atom in 3d space. It will, however, remove water and ions. The second array is of shape (atoms) and will contain the names for each atom as an mdtraj atom object. To actually read the names, it is good to cast to string. The last return is a frame object from mdtraj. It is returned just in case to allow vmd like selection language later on. As we do not plan to skip any frames or use a stride to skip frames, we do not need to provide these optional arguments. At this stage, we pretend to not know how many frames we
```

```
actually have available, so, even though it is costly, we want getList() to count the number of frames for us. Lastly, we supply a name, as getList() will do a backup save as a .npy numpy array."""

import ProteinPreprocessing.Preparation as Preparation

traj_1, names_1, chunk_1 = Preparation.getList(dcds=traj1_dcd, psf=traj1_psf, name="trajectory1")
traj_2, names_2, chunk_2 = Preparation.getList(dcds=traj2_dcd, psf=traj2_psf, name="trajectory2")

preparation = time.perf_counter()  # Time after preparation is done

"""Alignment: Protein structures are subject to internal motions during a simulation and can also rotate or even move as a whole. These motions will distort the later similarity measures. In order to only look at the internal motions and rearrangements, the protein structures need to be aligned individually and relative to each other. This alignment is done by Alignment.Align_states(), which returns two numpy arrays of the same form as Preparation.getList(). At the very least, Align_states takes both trajectory arrays and both name arrays, as returned by getList(). We set an accuracy of 0.03, so residues are considered, if their RMSF is lower than 3nm. We, once more, only need the standard input and can take the default for the rest. We have a small data set, so we can set Workers to the default of 1 for the RMSF calculation. No need to parallalize here. Lastly, we do not want to align according to a specific selection or want to have a specifially good alignment, so we can also keep the RMSD threshold as default."""


import ProteinPreprocessing.Alignment as Alignment

```

```
traj1_aligned, traj2_aligned = Alignment.Align_states(traj_1, traj_2,
    names_1, names_2, accuracy=0.05)

alignment = time.perf_counter()  # Time after alignment is done

"""Now, we make sure, that the trajectories have the right form and right
    length for the actual measures. The measures themselves, as they are
    made for each residue take numpy arrays of shape (atoms, timestep, 3)
    as opposed to the getList() or Alignment output of shape (timestep,
    atoms, 3). Additionally, we get another chance to input a Stride value
    to limit our dataset for test purposes, for instance. We can also set a
     number of frames, if we want to make sure, that both trajectories have
     the same length. As we continue with the changed data only, we can
    overwrite our old aligned numpy array with the transposed ones. We
    start out by preparing the selection on which we really want to compute
     our measures. Here, we chose to look at the CA atom in each amino
    acids backbone."""

traj1_index = Alignment.get_limited_selection_list(names_1, ["CA"])
traj2_index = Alignment.get_limited_selection_list(names_2, ["CA"])

"""Now we got all the inputs for SiMBols and we can create two Trajectory
    Objects. These Objects are still in the wrong form, so they need to be
    adjusted using the otc method (objects, time, coordinates). At the
    moment the array is still sorted as (time, objects, coordinates). We
    are also providing an index as calculated above, so SiMBols knows,
    whether to consider all objects."""

traj1_aligned = SiMBols.Trajectory(array=traj1_aligned, index=traj1_index)
traj2_aligned = SiMBols.Trajectory(array=traj2_aligned, index=traj2_index)



```

```python
traj1_aligned.otc(transpose=True)
traj2_aligned.otc(transpose=True)


"""Now we are ready to calculate the first distance measures. We supply
    the two trajectories to the measures and additionally supply a number
    of Workers for parallalized tasks to allow a quicker calculation. In
    this example, we will calculate the Wasserstein distance (wd) and the
    Frechet distnace (dfd).
We start by creating a comparer and give the two trajectories to it. The
    Comparer class has some built in functions to check the sanity of the
    input. We will forgo this check in the example"""

comp = SiMBols.Comparer(traj1_aligned, traj2_aligned)
comp.make_Check()
comp.cut()

print("And so it begins...")
Start = time.perf_counter()

comp.dfd(Workers=4)  # Calcuate the frechet distance on 4 processes
comp.wd()  # Calculate the Wasserstein distance

comp.save_all() # Save the results as numpy arrays

distances = time.perf_counter()  # Time after measures are done is done

print(f"Preparation: {round(preparation - start, 2)} second(s)!")
print(f"Alignment: {round(alignment - preparation, 2)} second(s)!")
print(f"Distances: {round(distances - alignment, 2)} second(s)!")
print(f"Total: {round(distances - start, 2)} second(s)!")

print("End Example Output")
```

(28) MatLab Version 2022a. 2022; Natick, Massachusetts: The MathWorks Inc.
